# Mapping the Antibody Repertoires in Ferrets with Repeated Influenza A/H3 Infections: Is Original Antigenic Sin Really “Sinful”?

**DOI:** 10.1101/2020.10.22.351338

**Authors:** Tal Einav, Martina Kosikova, Peter Radvak, Yuan-Chia Kuo, Hyung Joon Kwon, Hang Xie

## Abstract

The influenza-specific antibody repertoire is continuously reshaped by infection and vaccination. The host immune response to contemporary viruses can be redirected to preferentially boost antibodies specific for viruses encountered early in life, a phenomenon called original antigenic sin (OAS) that is suggested to be responsible for diminished vaccine effectiveness after repeated vaccination. In this study, we used a new computational tool called Neutralization Map to determine the hemagglutination inhibition profiles of individual antibodies within ferret antisera elicited by repeated influenza A/H3 infections. Our results suggest that repeated infections continuously reshape the ferret antibody repertoire, but that a broadly neutralizing antibody signature can nevertheless be induced irrespective of OAS. Overall, our study offers a new way to visualize how immune history shapes individual antibodies within a repertoire, which may help inform future vaccine design.

## Introduction

Rapidly evolving pathogens such as influenza frequently change their antigenicity in order to escape the host immune system, and the emergence of antigenically drifted strains necessitates the annual update of seasonal influenza vaccine components. Despite efforts to forecast which strain(s) will be most prevalent, a suboptimal or mismatched vaccine strain may occasionally be selected for vaccine production, resulting in reduced protection.^1-4^ In the US, influenza vaccine effectiveness in the past decades has fluctuated significantly from 10% in the 2004-2005 season^1^ to 60% in the 2010-2011 season (https://www.cdc.gov/flu/vaccines-work/effectiveness-studies.htm).^5^ While vaccine mismatch directly accounts for this low efficacy, pre-existing host immunity also influences vaccine performance.^3,6-12^

An individual’s exposure history, acquired through recurrent infections and/or vaccinations, shapes their unique antibody repertoire and influences their response to newly emerging influenza viruses.^6,10-19^ For example, several recent studies have reported that vaccine effectiveness is negatively correlated with vaccination frequency, with lower efficacy seen in more frequent vaccinees.^1,20-25^ While the exact mechanisms remain unknown, a suggested confounding factor is original antigenic sin (OAS) – a phenomenon where immune memory is recalled toward strains encountered early in life rather than to evolved viruses.^26 9,25,27-29^ On the other hand, residual antibodies from prior exposures may grant subsequent protection against viruses with similar antigenicity.^15-17,29^ These reports provide a glimpse of the complex interplay between prior and current immunity, highlighting the influence of immune imprinting that must be addressed in the field of vaccinology.

In this work, we profiled the hemagglutination inhibition (HAI) responses in ferrets after repeated influenza A/H3 infections and mapped the HAI antibodies elicited using a computational tool called Neutralization Map that characterizes antibody inhibition patterns.^30^ By tracking the progression of HAI antibodies following a series of infections, we demonstrated that repeated exposure to influenza can guide antibodies towards specific inhibition profiles irrespective of OAS.

## Results

### Sequential infections extended antibody cross-reactivity and induced broadly-neutralizing antibodies

We first conducted a sequential infection experiment in which seronegative ferrets were exposed to four influenza A/H3 viruses: A/Uruguay/716/2007 (Uruguay 2007, denoted as V_1_ throughout this work), A/Texas/50/2012 (Texas 2012, V_2_), A/Switzerland/9715293/2013 (Switzerland 2013, V_3_), and A/Hong Kong/4801/2014 (Hong Kong 2014, V_4_) as previously reported.^6^ We tracked the progression of antibodies developed after infection with V_1_ alone, followed by infection with the second (V_1_→V_2_), third (V_1_→V_2_→V_3_), and fourth virus (V_1_→V_2_→V_3_→V_4_), to demonstrate how the antibody repertoire was shaped by recurring exposures. As shown in Figure 1A, Uruguay 2007 (V_1_) infection elicited V_1_-specific ferret HAI titers with limited cross-reactivity towards viruses that emerged before 2005 or after 2007. Following each sequential infection with V_2_, V_3_, and V_4_, the cross-reactivity of ferret antisera gradually extended with geometric mean titers (GMTs) ≥ 80 against all A/H3 viruses in the panel except A/Philippines/2/1982 (Philippine 1982) that had disappeared from circulation more than three decades earlier (Figure 1B-1D).

**Figure 1.**
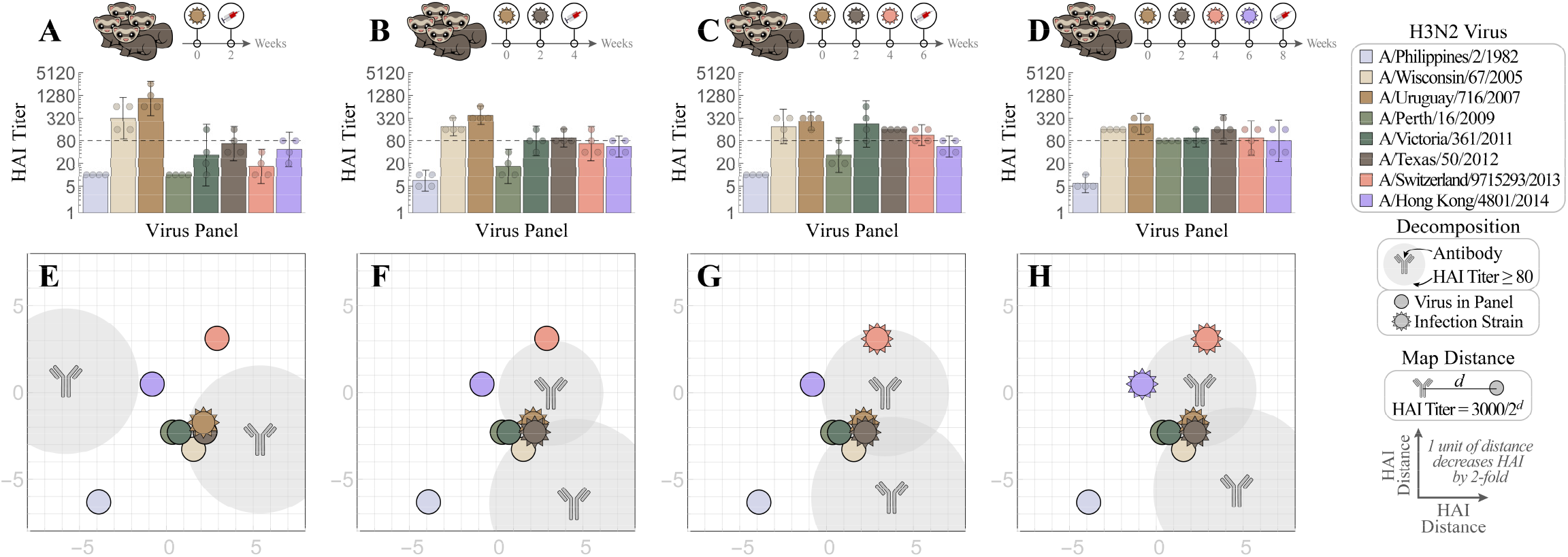
Tracking individual antibodies using hemagglutination inhibition (HAI) responses in ferrets during each stage of four sequential H3N2 infections. (A-D) Naïve ferrets were sequentially infected with V_1_=Uruguay 2007 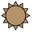, V_2_=Texas 2012 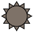, V_3_=Switzerland 2013 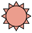, and V_4_=Hong Kong 2014 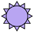. HAI titers are shown for each step of the infection: (A) V_1_, (B) V_1_→V_2_, (C) V_1_→V_2_→V_3_, and (D) V_1_→V_2_→V_3_→V_4_. Individual HAI titers are presented from four ferrets (points) with geometric means (bar graphs) and 95% confidential intervals (error bars). (E-H) These HAI measurements were decomposed to determine the individual antibodies elicited after infection with (E) V_1_ followed by (F) V_2_, (G) V_3_, and (H) V_4_. Each antibody signature (gray) is predicted to have an HAI titers ≥ 80 against any virus within the gray circle, with the size of this circle proportional to the fractional composition of the antibody in the serum [SI Methods]. An antibody-virus distance *d* denotes an HAI titer of 3000/2^*d*^. Virus coordinates were previously determined using a panel of monoclonal antibodies.^30^

To track antibody development throughout these four infections, we decomposed the HAI GMTs (Figure 1A-1D) using neutralization maps (Figure 1E-1H). On these maps, the positions of the eight influenza A/H3 viruses (identified by virus icons and solid dots) were previously determined using neutralizing data from a panel of monoclonal antibodies.^30^ Using these virus coordinates, the ferret antisera were computationally dissected to determine the number and location of antibody signatures that best match ferret HAI titers [SI Methods and Figures S1 and S2]. Each antibody signature may represent an amalgam of antibodies with similar inhibition profiles, although sufficiently distinct antibodies are decomposed as separate antibody signatures. The neutralization maps only show the strongest detectable antibody signatures that likely overwhelm the inhibition from weaker antibodies placed further away on the map. The Euclidean distance *d* between an antibody signature and a virus on the map translates into an HAI titer of 3000/2^*d*^, with potent antibodies lying near viruses against which they have a high titer. The gray regions surrounding each antibody signature indicate an HAI titer ≥ 80 against any virus that lies within (and we refer to such viruses as strongly inhibited).

As shown in Figure 1E, infection by V_1_ (Uruguay 2007) resulted in one “specific” antibody signature that strongly inhibited V_1_ and two nearby viruses – A/Wisconsin/67/2005 (Wisconsin 2005) and Texas 2012 – as well as another “non-specific” antibody signature that weakly inhibited all A/H3 viruses in the testing panel. With each subsequent infection (V_1_→V_2_, V_1_→V_2_→V_3_, and V_1_→V_2_→V_3_→V_4_), the specific antibody moved by 1-2 units in a manner that kept all infection strains strongly inhibited (within the gray circular regions surrounding each antibody Figure 1E-1H). Moreover, after the fourth infection, the GMTs across the entire virus panel were within 4-fold of one another (except for the older Philippines 1982 strain), indicating extended cross-reactivity of ferret antisera (Figure 1B-1D). While these maps depicted the average response of four ferrets, the individual maps of ferrets #1-4 in this cohort showed the same antibody trajectories (Figure S3). These progressional maps collectively suggest that a broadly-neutralizing antibody signature, which we define as an antibody with HAI titer ≥ 80 against all infection strains, can be guided into place by sequential exposures.

### Prior influenza exposures resulted in OAS and changed the inhibition profiles of elicited antibodies

We next compared the HAI response of ferrets infected with V_4_ alone with the responses elicited after one (V_3_→V_4_), two (V_2_→V_3_→V_4_), or three prior infections (V_1_→V_2_→V_3_→V_4_) to assess how exposure history affected the HAI antibody response to the final infection by V_4_. As shown in Figure 2A, infection by Hong Kong 2014 (V_4_) alone elicited higher HAI titers toward itself than to the other viruses in the panel. With additional prior exposures, ferret antisera always exhibited lower HAI GMTs toward V_4_ than to any of the earlier infection strains (Figure 2B-2D): for example, the V_3_→V_4_ infections elicited an HAI GMT of 34 toward V_4_ compared to the GMT of 135 toward V_3_, a typical OAS response that was also seen in Figure 1B-1D.

**Figure 2.**
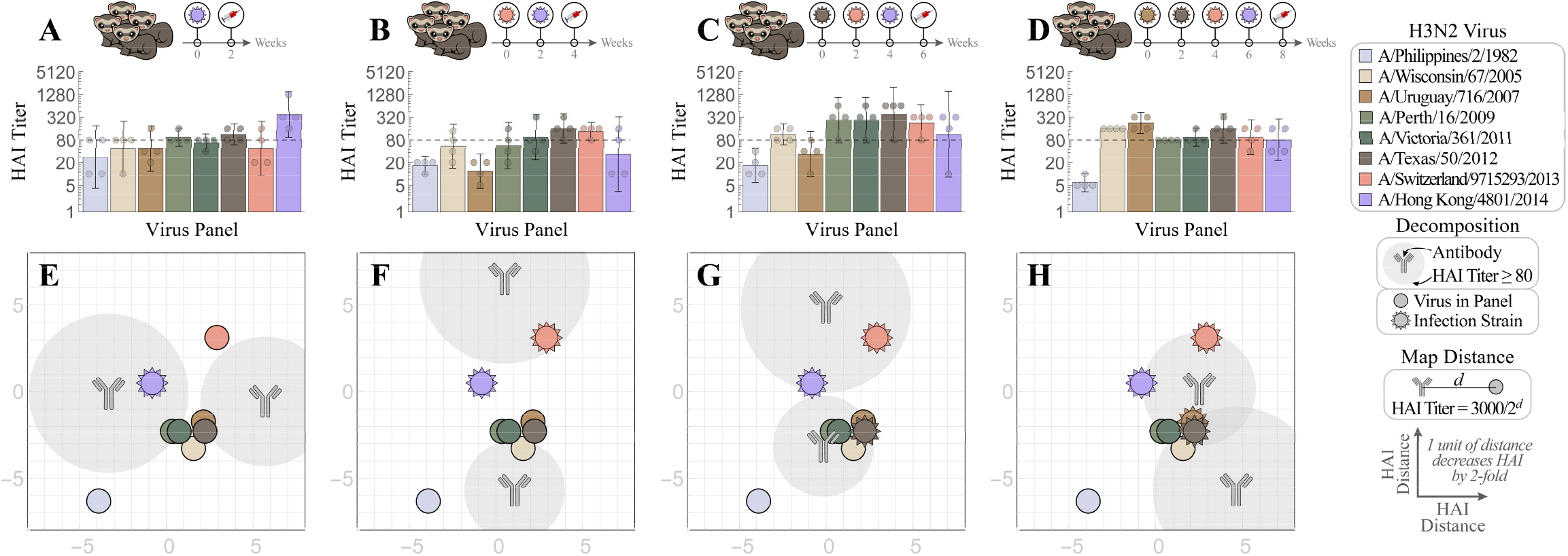
Mapping how exposure history shapes the ferret antibody response. Naïve ferrets were infected with (A) V_4_=Hong Kong 2014 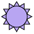, or with prior exposure to (B) V_3_=Switzerland 2013 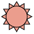, (C) V_2_=Texas 2012 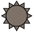, and (D) V_1_=Uruguay 2007 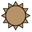. All antisera were analyzed via HAI after the final infection with Hong Kong 2014. Individual HAI titers are shown from four ferrets (points) and geometric means (bar graphs) with 95% confidential intervals (error bars). (E-H) Each set of HAI measurements was decomposed to determine the number of antibodies and their inhibition profiles. Each antibody signature (gray) is predicted to have an HAI titers ≥ 80 against any virus within the gray circle, with the size of this circle proportional to the fractional composition of the antibody in the serum [SI Methods]. An antibody-virus distance *d* denotes an HAI titer of 3000/2^*d*^.

To discern how prior exposures impacted subsequent antibody development, we decomposed the HAI GMTs on the neutralization maps (average response shown in Figure 2E-2H and individual responses shown in Figure S4). Upon infection by V_4_ alone, one specific antibody signature emerged that strongly inhibited V_4_ along with a non-specific antibody signature that displayed weak inhibition against the infection strain (Figure 2E). This same pattern was also observed in ferrets singly infected by V_1_, V_2_ or V_3_ (Figures 1-3 and S5). With each additional prior infection, OAS could be seen on the maps by noting that V_4_ lay further from the center of the gray antibody circles than the earlier infection strains, indicating a lower HAI response (Figure 2G-2H). Nevertheless, following the V_1_→V_2_→V_3_→V_4_ infections, ferrets developed antibodies that inhibited not only all four infection strains but also the other H3 viruses in the panel except Philippines 1982. Taken together, these results suggest that prior exposures affect the antibody response, but that a broadly-neutralizing antibody signature can be induced by repeated exposures, even in the presence of OAS.

**Figure 3.**
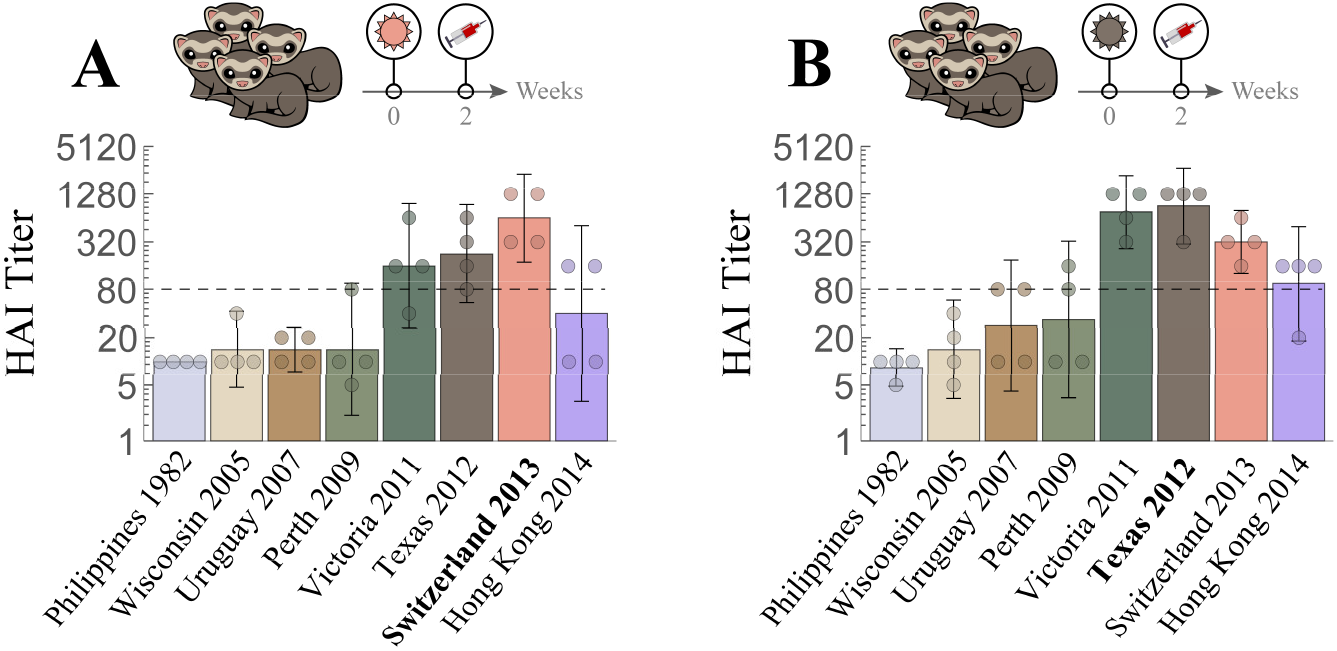
Cross-reactive hemagglutination inhibition (HAI) responses in ferrets with single H3N2 infection. Naïve ferrets were singly infected with either (A) V_3_=Switzerland 2013 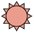 or (B) V_2_=Texas 2012 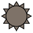. Individual HAI titers from four ferrets (points) and geometric means (bar graphs) are shown with 95% confidential intervals (error bars).

### The antibody profile elicited by sequential infections is strain-specific

We noticed that infection by V_1_→V_2_ resulted in potent inhibition (HAI ≥ 80) against both V_1_ and V_2_ (Figure 1F), whereas infection by V_3_→V_4_ resulted in antibodies that strongly inhibited V_3_ but not V_4_ (Figure 2F). To delve more deeply into this discrepancy, we analyzed the individual response of each ferret in these cohorts. Infection by V_1_ alone generated homologous HAI titers ≥ 640, whereas V_1_→V_2_ led to HAI titers ≥ 80 against both infection strains (Figure 4A-4D). In contrast, while infection by V_3_ alone generated homologous HAI titers ≥ 320, the subsequent infection V_3_→V_4_ resulted in different HAI responses in Ferrets #5 and #6 versus Ferrets #7 and #8 against V_4_ (Figure 4E-4H). Interestingly, Ferrets #5 and #6 with higher V_4_ cross-reactive titers after the first infection (left column of Figure 4E-4F) developed lower HAI titers following V_3_→V_4_. This can be seen on the neutralization maps, where the purple V_4_ virus lies within the gray circles signifying strong inhibition following V_3_ infection but not following V_3_→V_4_ (Figure 4E-4F). In contrast, Ferrets #7 and #8 which started with lower V_4_ cross-reactive titers developed higher titers following V_3_→V_4_. On the maps, this correspond to V_4_ transitioning from being outside the gray regions to inside them following V_3_→V_4_ (Figure 4G-4H). These results show that while repeated infections can result in robust HAI antibody responses against all infecting strains (Ferrets #1-4, 7-8), more complex inhibition patterns may arise (Ferrets #5-6).

**Figure 4.**
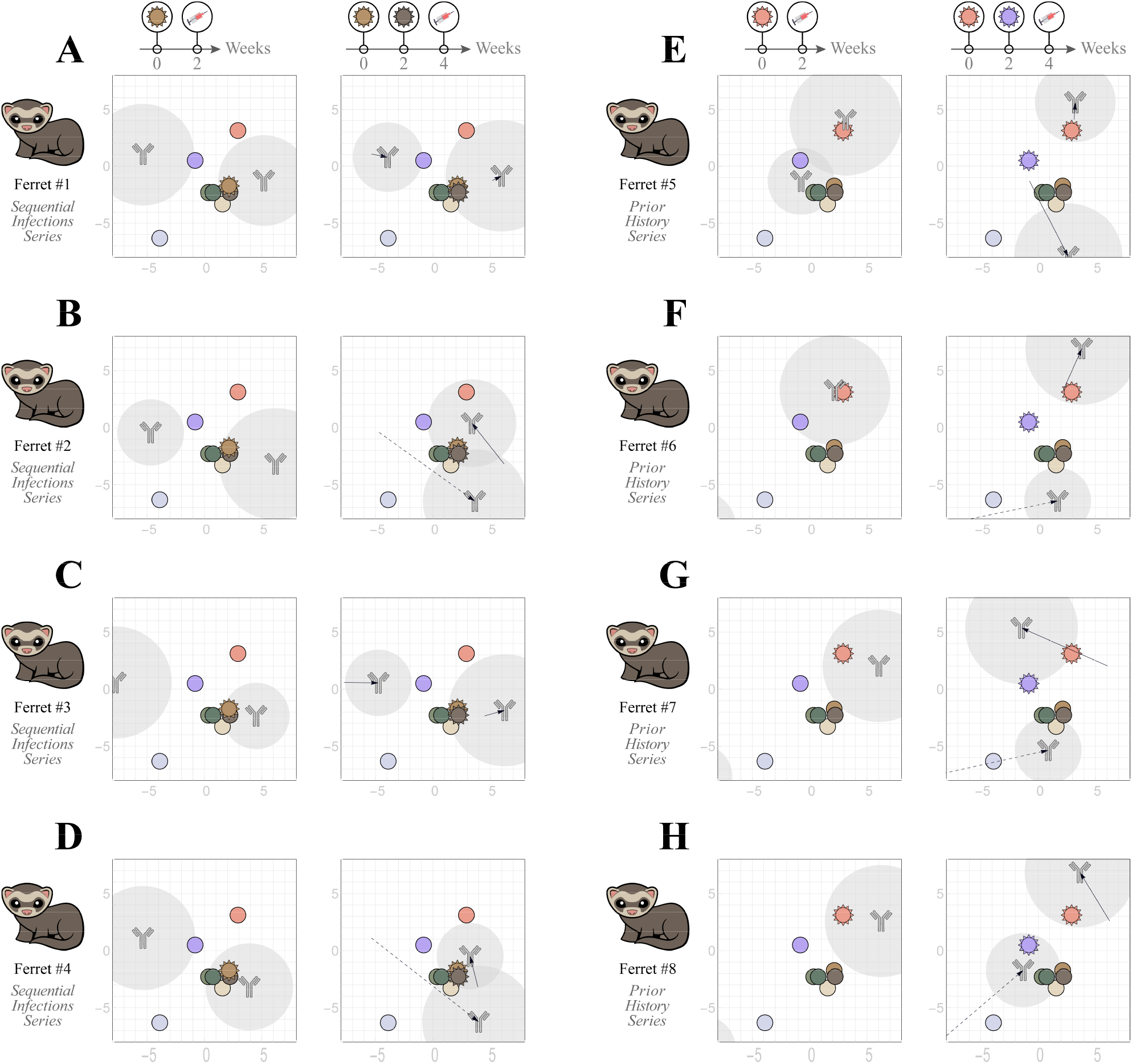
Individual ferret responses after two sequential infections. (A-D) Four ferrets were infected with V_1_=Uruguay 2007 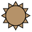 followed by V_2_=Texas 2012 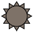. (E-H) Another group of four ferrets were infected with V_3_=Switzerland 2013 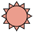 followed by V_4_=Hong Kong 2014 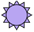. In each case, we decompose the antibody response after the first infection and track how each antibody changes after the second infection. Arrows with solid lines indicated the movement of an antibody signature along the map, with long dashed arrows denoting movement greater than 10 grid units that may represent multiple antibodies at the limits of detection [SI Methods].

## Discussion

It is estimated that most humans are infected with influenza by the age of 3 and continue to be reinfected by antigenically drifted strains every 5-10 years.^31,32^ Given the variability in infection histories and the stochastic processes involved in each specific infection, it is exceedingly difficult to determine the composition of pre-existing immunity and how it affects an individual’s antibody repertoire. In this study, we combined a ferret reinfection model with newly developed Neutralization Map to dissect the collective antibody response and characterize the properties of the constituent antibodies within.^30^ Using four recent H3N2 vaccine strains – Uruguay 2007 (V_1_) from the 2008-2010 seasons, Texas 2012 (V_2_) from 2013-2015, Switzerland 2013 (V_3_) from 2015-2016, and Hong Kong 2014 (V_4_) from 2016-2018 – we tracked the antibody footprints through each step of the sequential infections V_1_→V_2_→V_3_→V_4_ and deciphered the influence of prior exposures on the antibody response under four scenarios (V_4_ alone, V_3_→V_4_, V_2_→V_3_→V_4_, and V_1_→V_2_→V_3_→V_4_).

For each infection scheme, we found that the antibody repertoire contains at least one “specific” antibody signature that strongly inhibited the homologous virus and at least one “non-specific” antibody signature that weakly interacted with other H3 viruses in the panel. Along each step of the exposures V_1_→V_2_→V_3_→V_4_, both the specific and non-specific signatures tended to move closer to the infection strains. Eventually, a single cross-reactive antibody signature emerged that potently inhibited all four infection strains and the rest of the virus panel except Philippines 1982 (Figure 1H). These results likely reflect that with each infection, antibodies are refocused on conserved epitopes or structural regions across different infection strains.^33-35^

In a classical immune response to the same antigen, the reaction from the primary exposure is greatly magnified in subsequent encounters. OAS occurs upon exposure to antigenically-related viruses, where the response is skewed more heavily towards the earlier infection strains than to the latest strain. While this OAS phenomenon has been suggested to negatively affect vaccine effectiveness,^9,18,27,28,36,37^ seasonal vaccination can persistently extend the number of strains that the human antibody repertoire potently inhibits even when antibodies against earlier viruses are back-boosted.^38^ In this study, we observed OAS with each subsequent infection, regardless of the total number of exposures (Figures 1B-1D and 2B-2D). Despite the ubiquity of OAS, the crossreactivity of ferret antisera increased with each additional infection. These results build upon previous work which suggests that repeated exposures enhance antibody avidity.^6^

In general, repeated exposures resulted in antisera that strongly inhibited all infection strains. The sole exception was that 2 out of 4 ferrets had high V_4_ cross-reactive HAI titers in the preceding infection (with V_3_ alone, see left column of Figure 4E-4F) but exhibited a very weak HAI response towards V_4_ after infection by V_3_→V_4_. In contrast, the other two ferrets in this cohort as well as all four ferrets infected by V_1_→V_2_, elicited the typical response where both infection strains were strongly inhibited (Figure 4A-4D, 4G-4H). While it is unclear whether biological variation drives these observed differences, these results demonstrate that sequential infection can alter the patterns of an antibody response.

One feature of the neutralization maps is that the positions of the eight viruses in the panel were determined using data for human monoclonal antibodies, which may differ from ferret postinfection antisera. One such discrepancy is that the antigenic distance between V_1_ (Uruguay 2007) and V_2_ (Texas 2012) is small according to human monoclonal antibodies^39^ and postvaccination human sera,^3^ but that both viruses are considered antigenically distinct using ferret antisera raised from a single infection.^3^ In this study, substantially different HAI titers were observed across these two strains (Figures 1A-1B, 2A-2C, 3A-3B), which confirms that the ferret immune system treats V_1_ and V_2_ as antigenically distinct. Nevertheless, the full suite of HAI titers presented on the maps shows an average 2-fold error to the experimental measurements (Figures S1, S2C), demonstrating that the antigenic relationships among the majority of the viruses in the panel are the same across humans and ferrets.

In summary, by tracking the changes in the inhibition profile of ferret antisera induced by repeated influenza A/H3 infections, we demonstrated that an antibody could be guided along the map after a series of infections and that prior immune history can heavily influence the ferret antibody response. Despite the presence of OAS in each ferret infected by two or more viruses, a broadly neutralizing antibody signature that potently inhibited all infection strains was nevertheless produced (Figure 1H). While our current work was focused on HA head-specific antibodies, it does not consider antibodies directed towards the HA stem and neuraminidase that have also been shown to exhibit OAS and may influence the dynamics of this system.^40,41^ Future work that refines these antibody trajectories across multiple infections and multiple regions of an influenza virus may facilitate the development of more effective influenza vaccines.

## Methods

### Viruses

The panel of H3N2 viruses used for the study included A/Philippines/2/1982 (Philippine 1982), A/Wisconsin/67/2005 (Wisconsin 2005), A/Uruguay/10/2007 (Uruguay 2007), A/Perth/16/2009 (Perth 2009), A/Victoria/361/2011 (Victoria 2011), A/Texas/50/2012 (Texas 2012), A/Switzerland/9715293/2013 (Switzerland 2013) and A/Hong Kong/4801/2014 (Hong Kong 2014), each of which has served as the prototype for the H3N2 seasonal influenza vaccine component in the past decades. All H3N2 viruses were propagated in 9-10 days old embryonated eggs and aliquots were stored at −80°C until use.

### Ferret Infection

Seronegative male ferrets (Triple F Farm) at 15-16 weeks old were infected intranasally at two-week intervals with each of the four H3N2 viruses (V_1_=A/Uruguay/10/2007, V_2_=A/Texas/50/2012, V_3_=A/Switzerland/9715293/2013, and V_4_=A/Hong Kong/4801/2014) as previously reported.^6^ Ferrets were bled via venipuncture of the cranial vena cava at 14 days after each infection. Sera from four ferrets in each infection scheme were collected for HAI titer determination. All procedures were carried out in accordance with a protocol approved by the Institutional Animal Care and Use Committee of the Center for Biologics Evaluation and Research, US Food and Drug Administration.

### HAI Assay

Following pre-treatment with a receptor-destroying enzyme (Denka-Seiken), individual ferret sera were 2-fold serially diluted and were 1:1 (v/v) incubated with testing virus solution containing 4 hemagglutinin (HA) units per 25 μL at room temperature for 30 min before the addition of 50 μL of 0.75% guinea pig erythrocytes in the presence of 20 nM oseltamivir as previously described.^6,14^ The endpoint HAI titer was defined as the reciprocal of the highest serum dilution that yielded a complete HA inhibition, and a titer 5 was assigned if no inhibition was observed at the starting 1:10 serum dilution. HAI geometric mean titers (GMTs) were calculated along with the 95% confidence intervals.

### Decomposing Ferret Antisera on the Neutralization Maps

The positions of the eight H3N2 viruses in the testing panel were previously determined using neutralizing titers from 6 human monoclonal antibodies.^30^ These maps quantify how an individual antibody simultaneously inhibits all eight H3N2 viruses, where an antibody-virus distance *d* on the map corresponds to an HAI titer of 3000/2^*d*^. A key assumption of this analysis is that each antibody can be represented by a single point on the map, so that scanning the map exhaustively searches through the possible inhibition profiles for any antibody targeting the head domain of influenza hemagglutinin.

To decompose each serum, we searched through an increasing number of antibodies to determine which combination of coordinates and stoichiometries best matched the HAI titers of the eight H3N2 viruses in the panel [SI Methods]. The process ended when adding an additional antibody did not substantially improve the match with the experimental measurements. We validated the resulting decompositions in two ways. First, we determined that the HAI titers presented on the maps were on average only 2-fold off from the experimental measurements, demonstrating that these maps can capture the trends in the HAI profiles (Figure S1). Next, we decomposed each antiserum using HAI titers from only four viruses (Philippines 1982, Wisconsin 2005, Switzerland 2013, Hong Kong 2014). The predicted titers against the remaining four viruses were only 5-fold off from the experimental measurements, comparable to the results from earlier work analyzing mixtures with known antibody composition (Figure S2).^30^

## Author contributions

H Xie conceived and designed the ferret study. M Kosikova, P Radvak, H Xie, and Y-C Kuo conducted the ferret infection experiments. M Kosikova, H Xie, P Radvak and H J Kwon performed the HAI assays. T Einav conceived and designed Neutralization Map. T Einav, M Kosikova and H Xie analyzed the HAI data. H Xie and T Einav wrote the paper.

## Disclaimer

The findings and conclusions in this article have not been formally disseminated by US Food and Drug Administration and should not be construed to represent any Agency determination or policy.

