## Supplementary Information for "Mapping the Antibody Repertoires in Ferrets with Repeated Influenza A/H3 Infections: Is Original Antigenic Sin Really “Sinful”?"

### Determining the Virus Positions on the Neutralization Maps

The positions of the eight H3N2 viruses in the panel had been previously determined using neutralization data from 6 human monoclonal antibodies targeting the head of influenza hemagglutinin (HA).[1] 50% inhibitory concentrations ($IC_{50}$s) were determined between each antibody and virus pair, and their positions on the Neutralization Maps were fixed using 2D multidimensional scaling on the $\log_{10}(IC_{50})$ values.

The proximity between two viruses determines their antigenic similarity. For example, two strains that lie near one another on the map are expected to be similarly neutralized by any monoclonal antibody (and hence by any serum). We assume that the neutralization and hemagglutination inhibition (HAI) for HA head-targeting antibodies is identical, which is reasonable if neutralization is mediated by an antibody occluding HA binding to the target cells.

Compared to the original maps presented in Einav *et al.*[1], we made several cosmetic changes to tailor these maps towards HAI data. First, whereas 1 grid unit on the original neutralization maps represented a 10-fold decrease in neutralization, we decreased the spacing between gridlines by a factor of $\log_{10}(½)$=3.3 so that 1 grid unit represents a 2-fold decrease in this manuscript. Second, we translated all virus coordinates (by 5.5 units to the right and 3 units down) in order to center the viruses on the map, since it is only relative distances (instead of absolute coordinates) that matter on the map. Neither of these changes affect the resulting serum decompositions in any way.

We also note one minor discrepancy between the virus strains in our panel versus the viruses used in the original work.[1] In the original work, A/Brisbane/10/2007 was included in the virus panel.[1] For this project, we infected ferrets and measured their response against the A/Uruguay/716/2007X175C strain (denoted by $V_1$) which is antigenically A/Brisbane/10/2007-like. Hence, we used the map coordinates inferred for A/Brisbane/10/2007 from the original work as the coordinates for A/Uruguay/716/2007X175C in this work.

### Converting HAI serum dilution into absolute titers

As shown in Figure 1A-1D, Figure 2A-2D, and Figure 3, the ability of sera to inhibit the virus panel is measured in titers, that is, as the largest dilution of the serum capable of inhibiting hemagglutination. However, distance on the neutralization maps must be defined in absolute units in order to model how combinations of antibodies will collectively inhibit a virus. More precisely, an antibody-virus distance of $d$ units translates into $IC_{50}=(10^{-10}$ Molar$)(2^d)$, which represents the concentration of the antibody necessary to neutralize the virus by 50%.
As in previous work, we converted HAI titers into absolute Molar units by scanning across different conversion factors.[1] For each factor, we decomposed all of the ferret sera and quantified the error between the resulting HAI titers from the inferred antibodies and the measured titers. The conversion factor that minimized the error corresponded to an $IC_{50}=(3000\cdot 10^{-10}$ Molar)/(HAI titer), so that a map distance $d$ between an antibody and virus translates into an HAI titer$=3000/2^d$. Note that we used this same global conversion factor for all of the sera analyzed in this work.

### Determining the Antibody Positions on the Neutralization Maps

We decomposed each serum by determining which set of antibody coordinates and stoichiometries best match the HAI titers of this serum against our virus panel. As previous described, decomposition proceeds by considering $n$=1, 2, 3... antibodies until the error of the decomposition decreases below a set threshold.[1] We first compute the best decomposition using a single head antibody, and then assessed two head antibodies with any stoichiometry (although each antibody must comprise ≥20% of the mixture to avoid adding trivial antibodies that do not appreciably change the inhibition profile). As in previous work, we accepted the 2-antibody decomposition provided that the error between the decomposition's HAI titers and the measured titers decreased by at least 20% and above a noise threshold.[1] If the 2-antibody decomposition passed the threshold, we went on try all 3-anibody decompositions, and we continued this process until adding another antibody no longer met the threshold requirement for the decrease in error, at which point the algorithm ends (i.e. if the $n$-antibody decomposition does not pass either threshold, then we describe the mixture using the best ($n$-1)-antibody decomposition).

The abundance of each antibody in a mixture is represented by the size of the gray circle surrounding it. For any virus lying within a gray circle, the antibody at the center of that circle is predicted to have an HAI titers ≥ 80 against it. If an antibody comprises a fraction $f$ of a mixture, then its ability to inhibit a virus decreases by $f$-fold. Hence, while a monoclonal antibody would be surrounded by a circle of radius $r_{\text{mAb}}$=5.2 grid units (since $3000/2^{r_{\text{mAb}}}=80$), an antibody that comprises a fraction $f$ of the serum will be surrounded by a circle of radius $r_{\text{mAb}} - \log_2(1/f)$. For example, the specific antibody signature in Figure 1H that strongly inhibits all four infection strains was found to comprise 25% of the influenza repertoire while the non-specific signature comprises 75% of the repertoire. Hence the radii surrounding these two antibody signatures were 3.2 and 4.8 grid units, respectively.

When multiple antibodies are present in a serum, their collective inhibition or neutralization against a virus is computed using a competitive binding model,

$$IC_{50,\text{Competitive}} = \left(\sum_j f_j/IC_{50}^{(j)}\right)^{-1},$$

where $f_j$ represents the fractional composition of the $j^{\text{th}}$ antibody and $IC_{50}^{(j)}$ denotes the concentration at which this monoclonal antibody would neutralize the virus by 50%. In this way, any combination of points (which determines $IC_{50}^{(j)}$ through antibody-virus map distance) at any stoichiometry ($f_j$) can be translated into the mixture's collective HAI titer against the virus panel.

### Characterizing the Full Suite of Ferret Antisera

We first investigated how well the 2D maps can represent the antibody-virus inhibition profiles seen across the sera. Figure S1A shows the decomposition of the serum elicited at the end of our series of infections $V_1 \rightarrow V_2 \rightarrow V_3 \rightarrow V_4$, with the adjacent table showing the close correspondence between the measured HAI titers and the titers shown on the map. Figure S1B extends this analysis to all antisera considered in this work. Each point represents a pair of measured and map titers (like the eight pairs shown in Panel A), and the points are drawn with low opacity to show the trend towards the diagonal line, indicating that the map titers accurately correspond to the measured values (with a coefficient of determination $R^2$=0.7).

The decomposition shown in Figure S1A predicts that the serum elicited after all four infections contains an antibody that can single-handedly inhibit $V_1$, $V_2$, $V_3$, and $V_4$ with an HAI titer $\geq 80$ at its concentration in the serum. Although we do not experimentally dissect the serum and confirm the existence of such an antibody, we validate our decompositions by using a subset the HAI titers to predict the remaining titers across our virus panel. In other words, rather than performing the decomposition using the titers against all eight viruses (Figure S2A), we decompose this serum using the inhibition titers against four viruses (Philippines 1982, Wisconsin 2005, Switzerland 2013, Hong Kong 2014) and use the inferred antibody signatures to predict the serum inhibition against the rest of the panel (gray viruses in Figure S2B).

Carrying out this analysis for all antisera, we find that the predicted titers are 5.1-fold off from the measurements on average (Figure S2C), a notable achievement given that the measured HAI titers spanned a 500-fold range. As additional viruses are used for the decomposition, the antibody signatures are determined more precisely, and the prediction error will approach the 2.3-fold error achieved when using the full suite of data (Figure S2C).

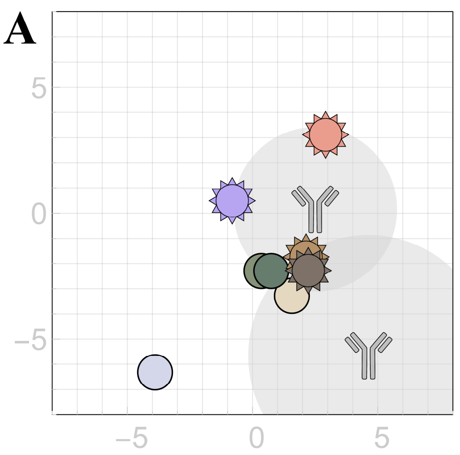


| Virus | Measured HAI | Map HAI |
|---|---|---|
| Philippines 1982 | 6 | 6 |
| Wisconsin 2005 | 160 | 154 |
| Uruguay 2007 | 226 | 206 |
| Perth 2009 | 80 | 85 |
| Victoria 2011 | 95 | 102 |
| Texas 2012 | 160 | 165 |
| Switzerland 2013 | 95 | 97 |
| Hong Kong 2014 | 80 | 76 |

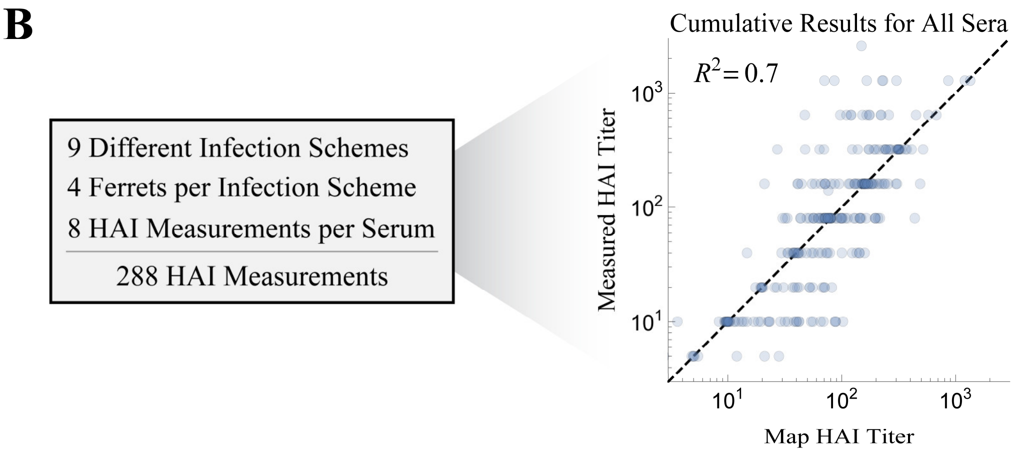


**Figure S1. Characterizing the accuracy of the two-dimensional neutralization maps.** (A) Serum decomposition mapping the antibody response after the four virus infections $V_1 \to V_2 \to V_3 \to V_4$. The table shows the measured HAI values together with the HAI values represented on the map by translating map distance $d$ into an HAI titer (given by $3000/2^d$ for each individual antibody, and the titers from both antibodies are combined using the competitive binding model as described in the SI Methods). (B) Comparing the measured and map HAI titers across all ferret antisera analyzed in this work.

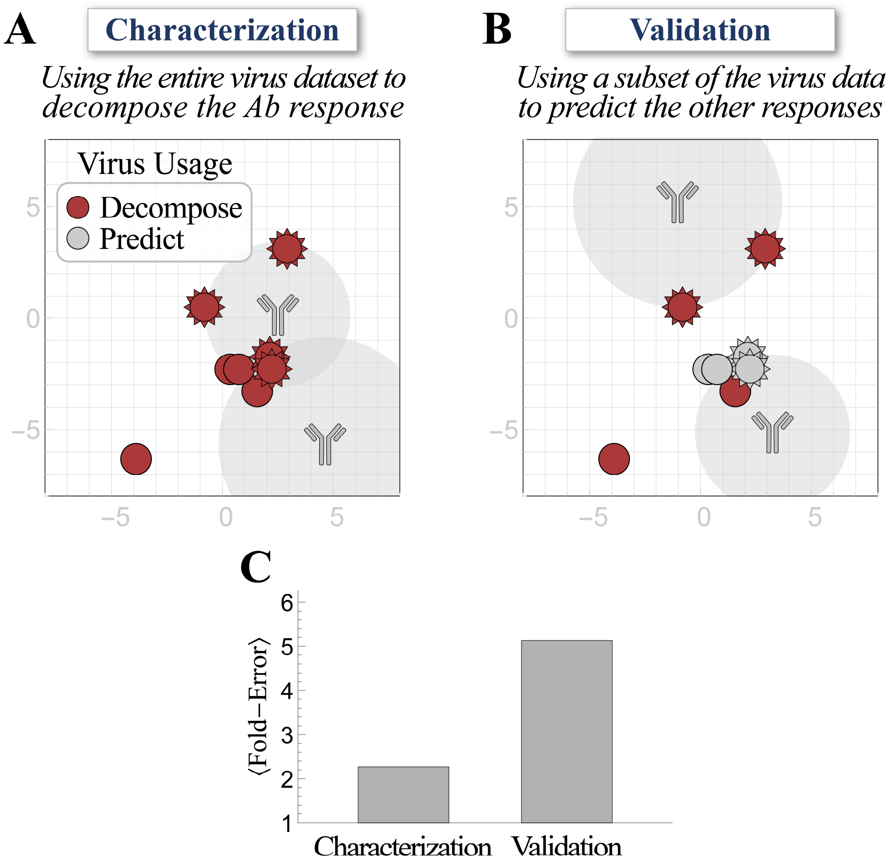


**Figure S2. Validating the antibody decompositions using a subset of data.** (A) All of the serum decompositions shown in this work were performed using HAI titers from the entire virus panel. (B) A decomposition may also be performed on a subset of the virus panel (red). The resulting antibody locations predict the HAI titers of the remaining viruses (gray) which can be compared with the measured values. (C) The average decomposition error from both methods.

### Decomposing the Individual Ferret Responses Across Repeated Infections

In Figure 1E-1H and Figure 2E-2H, we decomposed the average response (HAI GMTs) of four ferrets in each vaccination group. While this provides a compact representation of these four replicates, the inherent stochasticity of the immune response will generate different antibodies in each ferret, and averaging occludes this variability. Moreover, averaging titers can introduce artifacts in the data, since the geometric mean of titers produced by monoclonal antibodies may produce an inhibition profile that is impossible for a monoclonal antibody to achieve (and hence may result in a polyclonal decomposition).

Because of these considerations, Figure S3 shows the decompositions of each individual ferret antiserum shown in Figure 1. Note that the same four ferrets progressed through all four infections (i.e. Ferret #1 was infected with $V_1$, then 2 weeks later it was bled right before it was infected with $V_2$, and so on). Therefore, these decompositions enable us to track the evolution of these antibodies in every ferret following each exposure (arrows in Figure S3), providing a more nuanced perspective to quantify how infections reshape the antibody repertoire.

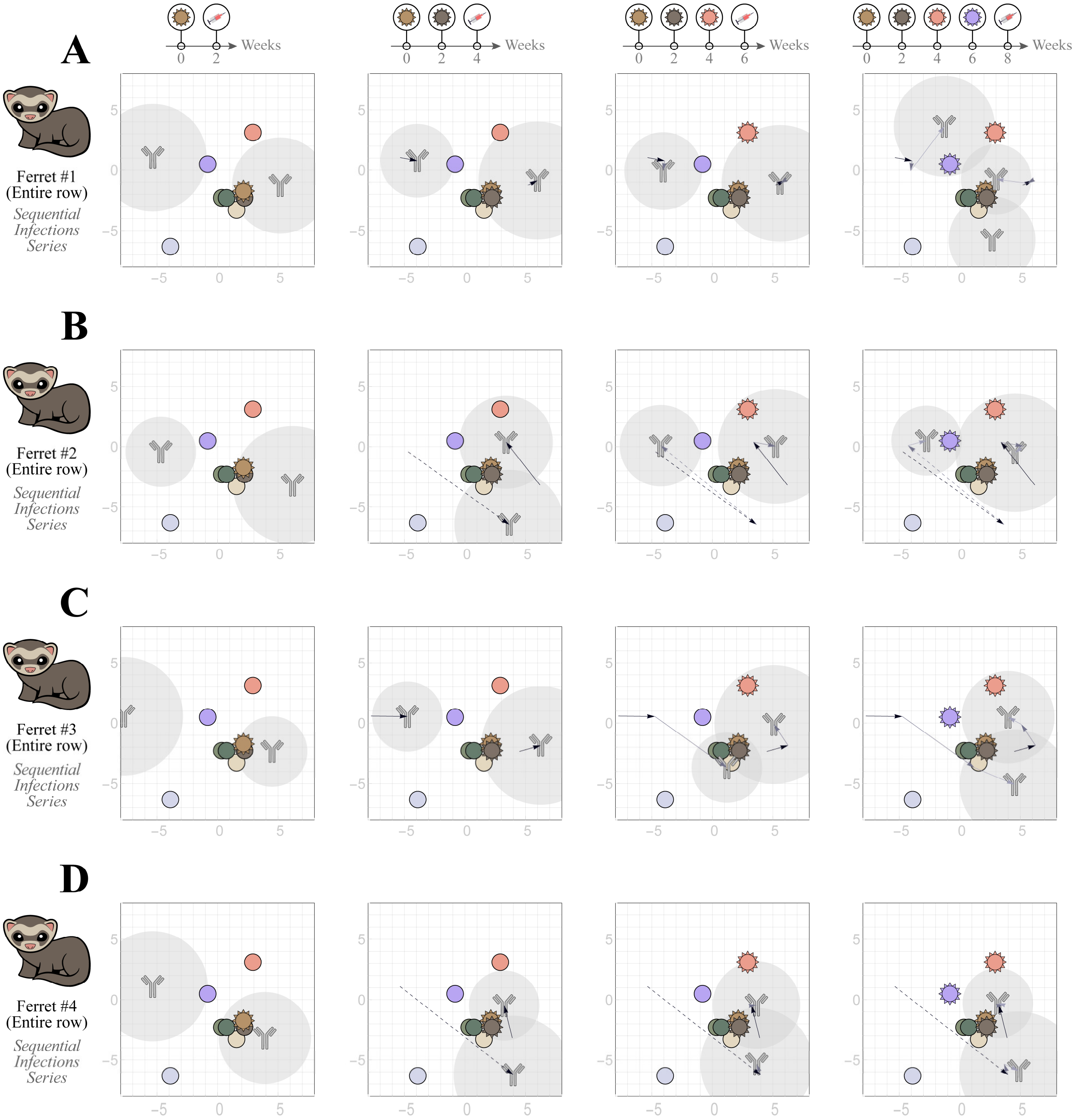


**Figure S3. Tracing individual ferret responses through four sequential infections.** Neutralization maps created using HAI titers from individual ferrets infected by (A) $V_1$=Uruguay 2007 followed by (B) $V_2$=Texas 2012, (C) $V_3$=Switzerland 2013, and lastly (D) $V_4$=Hong Kong 2014. Arrows track the motion of each antibody, with dashed arrows denoting movement greater than 10 grid units that may represent multiple antibodies at the limits of detection. The average response in Figure 1 depicts the geometric mean of the HAI titers shown in each column.

---

We note that the resolution of each decomposition is inherently limited by the distribution of viruses across the map. Each virus serves as a "detector" for the antibody nearest to it, and at least three measurements are needed to accurately pinpoint an antibody. Therefore, many of the antibody signatures in these sera may go undetected because their inhibition is overwhelmed by antibodies that lie nearer to the viruses. For example, the large movement seen in the non-specific antibody in Ferret #2 (dashed arrows in the two middle columns of Figure S3B) likely indicate that this ferret produced an antibody in the left region of the map and another antibody in the bottom-right, but there was only sufficient resolution to detect one of these antibodies each time. To distinguish these cases, we drew any arrow whose length was greater than 10 grid units using dashed lines to indicate that it may connect two different antibodies. As more decompositions are carried out on maps that contain more viruses, we will be able to determine how far an antibody can move with each infection as well as quantify the noise in this process.

Figure S4 decomposes the individual responses shown in Figure 2. Because the infection history differed each time, 16 different ferrets were used in this experiment. Therefore, we could not track the trajectories of the individual antibodies as in Figure S3. As noted in the main text, the greatest variability was seen in the responses towards $V_4$. While exposure to any of the other three infection strains mainly resulted in potent inhibition against them (i.e. in these viruses lying within the gray antibody circles), 6 out of 16 of these ferret responses showed $V_4$ outside of these gray regions, signifying a weaker response even after exposure to this strain.

### Decomposing the Individual Ferret Responses to Single Infections

To make contact with the standard characterization of viruses using ferret antisera raised from a single infection, Figure S5 shows the decompositions of sixteen ferrets exposed to one of the infection strains considered in this work. Each column shows the variability of the response across four ferrets exposed to the same strain. While there was notable similarity between the specific antibody signatures (closest to the infection strain), marked differences could be seen between the non-specific antibodies elicited. These results suggest that while a single infection may yield consistently large homologous titers, the resulting inhibition against antigenically distinct viruses may be highly varied. Moreover, these differences caused by the non-specific antibodies (e.g. whether infection by $V_3$ happens to elicit a non-specific antibody that potently inhibits $V_4$) may play an important role in shaping future responses (e.g. the response to subsequent infection by $V_4$), especially in the human immune repertoire where dozens of viruses may be encountered over a lifetime.

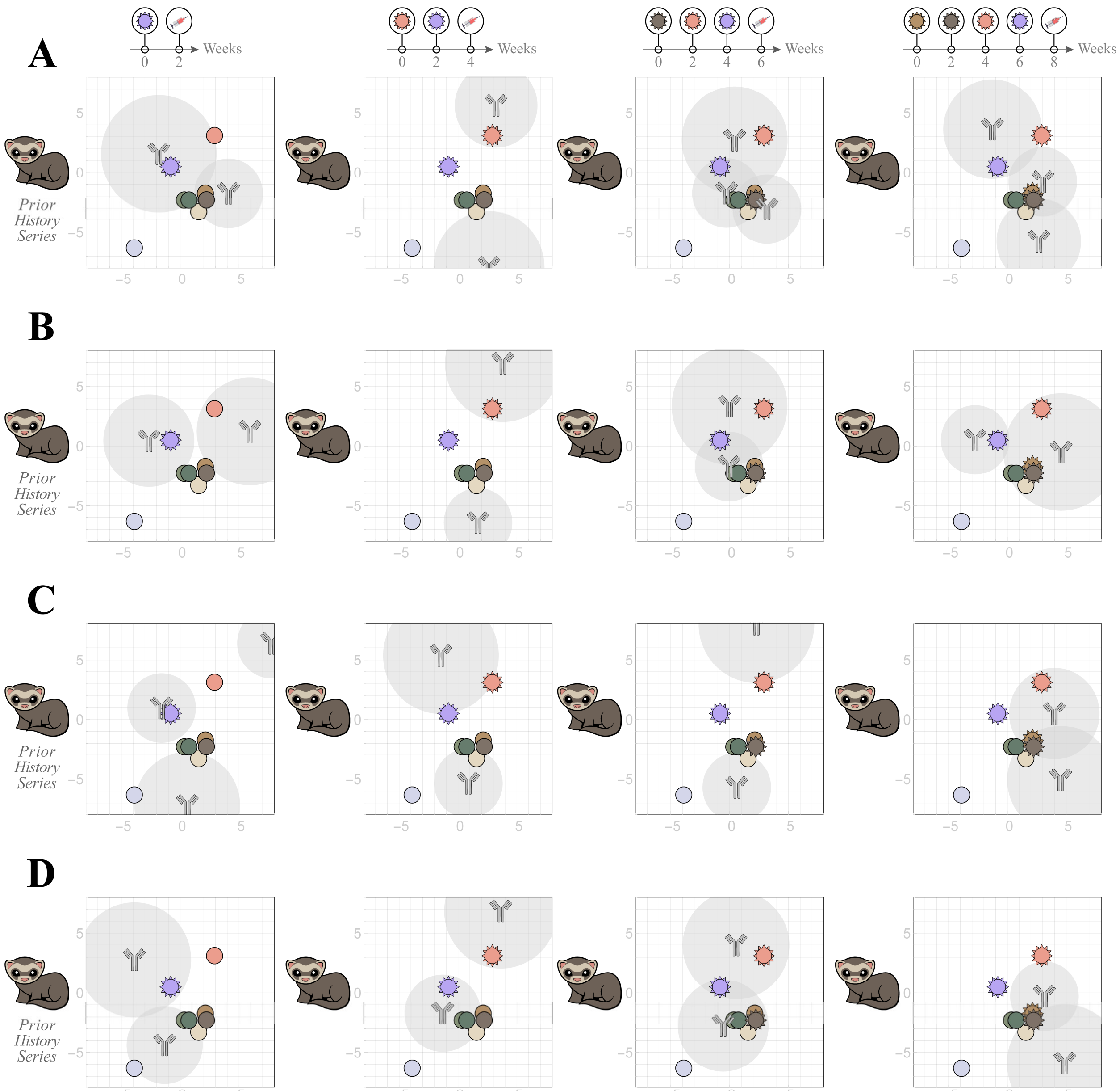


**Figure S4. Tracing individual ferret responses with four different prior infection histories.** Neutralization maps created using HAI titers from individual ferrets infected by (A) $V_4$ alone, (B) $V_3 \rightarrow V_4$, (C) $V_2 \rightarrow V_3 \rightarrow V_4$, or (D) $V_1 \rightarrow V_2 \rightarrow V_3 \rightarrow V_4$. A total of sixteen ferrets were tracked. The average response in Figure 2 depicts the geometric mean of the HAI titers shown in each column.

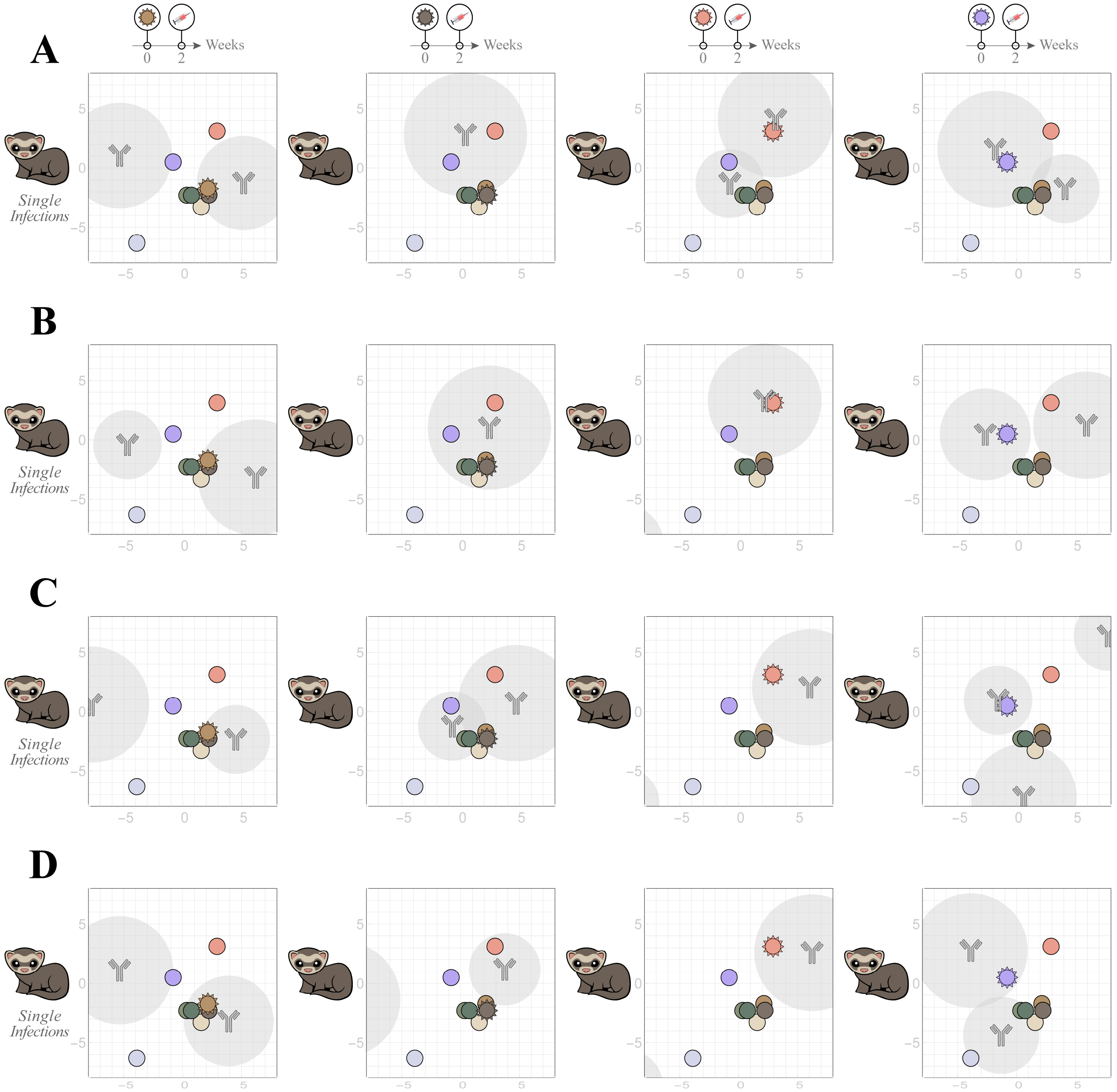


**Figure S5. Antibody repertoires from single infections of ferrets.** Neutralization maps created using HAI titers from individual ferrets infected by either (A) $V_1$, (B) $V_2$, (C) $V_3$, or (D) $V_4$ alone.
